# A Scoping Review of Mathematical Models Covering Alzheimer’s Disease Progression

**DOI:** 10.1101/2022.12.07.519498

**Authors:** Seyedadel Moravveji, Nicolas Doyon, Javad Mashreghi, Simon Duchesne

## Abstract

Alzheimer’s disease is a complex, multi-factorial and multi-parametric neurodegenerative etiology. Mathematical models can help understand such a complex problem by providing a way to explore and conceptualize principles, merging biological knowledge with experimental data into a model amenable to simulation and external validation, all without the need for extensive clinical trials.

We performed a scoping review of mathematical models of AD with a search strategy applied to the PubMed database which yielded 846 entries. After applying our exclusion criteria, only 17 studies remained from which we extracted data, focusing on three aspects of mathematical modeling: how authors addressed continuous time, how models were solved, and how the high dimensionality and non-linearity of models were managed. Most articles modeled AD at the cellular range of the disease process, operating on a short time scale (e.g., minutes; hours), i.e., the micro view (12/17); the rest considered regional or brain-level processes, with longer timescales (e.g., years, decades) (the macro view). Most papers were concerned primarily with *Aβ* (*n* = 8), few modeled with both *Aβ* and tau proteins (*n* = 3), and some considered more than these two factors in the model (*n* = 6). Models used partial differential equations (*PDEs*; *n* = 3), ordinary differential equations (*ODEs*; *n* = 7), both PDEs and ODEs (*n* = 3). Some didn’t specify the mathematical formalism (*n* = 4). Sensitivity analyses were performed in only a small number of papers (4/17).

Overall, we found that only two studies could be considered valid in terms of parameters and conclusions, and two more were partially valid. The majority (*n* = 13) either was invalid or there was insufficient information to ascertain their status. While mathematical models are powerful and useful tools for the study of AD, closer attention to reporting is necessary to gauge the quality of published studies to replicate or continue with their contributions.

## Introduction

Alzheimer’s disease (**AD**) is a neurodegenerative disorder that results in severely reduced cognition, loss of autonomy, eventual physical weakness, and death [22]. The occurrence of AD is strongly related to aging and the whole disease process covers a period estimated to last a minimum of 10-20 years before diagnosis [15]. This prodromal phase is thought to depend on physical, genetic, and epigenetic factors, all to be conclusively determined [15].

Indeed, AD is difficult to diagnose in its earlier phases, since its etiology remains unclear. While multiple hypotheses abound (e.g., beta-amyloid (*Aβ*) cascade hypothesis [10, 16, 19, 20]; tau protein hypothesis [17]; oligomer cascade hypothesis [19], none have been proven beyond doubt nor been translated to a positive clinical outcome after interventions. Rather, it is becoming clear in the community that AD is a multi-factorial disease, influenced by several risk factors, and hence also multi-parametric. Therefore, early detection of AD and identification of a cure continue to be major challenges for the scientific community given both this lack of etiological clarity and apparent complexity.

As a consequence, a fundamental obstacle of AD research becomes the sheer number of biological, environmental, and psychosocial variables that must be considered, and by extension the geometrically increasing number of interactions between variables. These become difficult, if not impossible to assess thoroughly via either clinical trial experiments, as the logistics of recruiting a sufficient large number of individuals to achieve reasonable statistical power become daunting, without mentioning the decades-long timeframe required to follow these individuals along this neurodegenerative process. Conversely, while preclinical models can often provide clarity with respect to specific variables and their interactions, they suffer from problems of generalizability to pseudo-sporadic AD in human populations.

Mathematical modeling on the other hand is a great tool to understand complex mechanisms such as AD. Mathematical models provide a way to explore and conceptualize principles by merging biological knowledge with experimental data into model simulations [6]. In models, the physical reality is abstracted into entities and parameters that can help us more easily understand relationships. In the case of AD, it could help us figure out causal mechanisms and therefore propose targets for disease prevention. Accurate models should help with intervention planning and trial, by performing *in sillico* experiments of effects and drug interventions. Models however are only as good as the assumptions on which they are based. Indeed, if a model makes predictions that are out of line with observed results, or that cannot be verified experimentally altogether, either the entity relationships must be modified, or initial assumptions must change.

To guide our group in the elaboration of a comprehensive, multi-factorial mathematical model of AD, we elected to perform a scoping review of this nascent literature, the results of which are presented in the following sections. We paid particular attention to three aspects of mathematical modeling: first, how authors addressed continuous time, as relevant in the case of AD; secondly, how models were solved, given these different time scales at which the interactions between variables operate; and finally, how were high dimensionality and non-linearity of models managed, up to and including parameter sensitivity.

## Methods

### Literature search strategy and sources

We performed a scoping review based on the PRISMA standards [24]. We searched the PubMed database for peer-reviewed, original research journal papers in English published through March 29, 2022. Our search terms were “Mathematical model,” “Alzheimer’s disease,” and “Not animal”.

### Study selection process

Using the Covidence systematic review system (Melbourne, Australia), we removed duplicate entries after the initial search, then did the first screen based on article titles and abstracts. We then completed a full-text evaluation of each article. Our goal was to identify multi-factorial mathematical models of AD, applied to a human population. Consequently, we disregarded approaches that were too narrow (e.g., only one protein) or animal-centered. Two impartial reviewers independently conducted this process (S.M. and S.D.), and consensus solved conflicts.

### Data extraction

We extracted the following characteristics from all included studies (cf. Table 1): Title, Lead author contact details, Country(ies) of origin of authors, Aim of study, Temporal scale (micro or macro) (see section 3.2), Multi-structure or not (see section 3.2), Validated or not, Interesting summary figure, Study funding sources, Possible conflicts of interest for study authors. Information regarding mathematical modelling was also extracted (cf. Table 2), including Model type (mathematically speaking); If ordinary differential equations (**ODE**), what solver was used; Given the solver, was it adapted to stiff equations; along with the explanation for each study, i.e., What concepts/entities were captured by the model.

**Table 1.** Studies characteristics.

|  |  | Title | Object | Time Scale |
| --- | --- | --- | --- | --- |
| 1 | Anastasio 2011 | Data-driven modeling of Alzheimer's disease pathogenesis. | $A\beta$ | Micro |
| 2 | Bertsch 2017 | Alzheimer's disease: a mathematical model for onset and progression. | $A\beta$ | Macro |
| 3 | Fornari 2019 | Prion-like spreading of Alzheimer's disease within the brain's connectome. | Tau protein | Micro |
| 4 | Han 2020 | Computational modeling of the effects of autophagy on amyloid-β peptide levels. | $A\beta$ | Micro |
| 5 | Han 2016 | A Theoretical Analysis of the Synergy of Amyloid and Tau in Alzheimer's Disease. | $A\beta$, Tau | Micro |
| 6 | Hao 2016 | Mathematical model on Alzheimer's disease. | $A\beta$, Tau, Astrocytes, Microglia, Macrophages, GSK3, NFT, APP, $TNF_{\alpha}$, IL-10, TGF, MCP-1, ROS, HMGB1 | Macro |
| 7 | Helal 2014 | Alzheimer's disease: analysis of a mathematical model incorporating the role of prions. | $A\beta$ oligomers, $A\beta$ plaques, and cellular prion proteins | Micro |
| 8 | Helal 2019 | Stability analysis of a steady state of a model describing Alzheimer's disease and interactions with prion proteins. | $A\beta$ peptides, $A\beta$ oligomer | Micro |
| 9 | Hoore 2020 | Mathematical Model Shows How Sleep May Affect Amyloid-β Fibrillization. | $A\beta$, $A\beta$ oligomer, microglia, CSF | Macro |
| 10 | Kuznetsov 2018 | How the formation of amyloid plaques and neurofibrillary tangles may be related: a mathematical modeling study. | Tau, APP external $A\beta$ plaques, intracellular tangles | Micro |
| 11 | Kuznetsov 2018 | Simulating the effect of the formation of amyloid plaques the on aggregation of tau protein. | $A\beta$, Tau, APP, neurofibrillary tangles | Micro |
| 12 | Lindstrom 2021 | From reaction kinetics to dementia: A simple dimer model of Alzheimer's disease etiology. | $A\beta$ Oligomer, $A\beta$ monomers, dimers, trimers | Macro |
| 13 | Liu 2015 | Evaluating Alzheimer's Disease Progression by Modeling Crosstalk Network Disruption. | $A\beta$, $A\beta$ Oligomer, P-Tau, Tau, MCI, GSK3 | Micro |
| 14 | Pallitto 2001 | A mathematical model of the kinetics of beta-amyloid fibril growth from the denatured state. | $A\beta$ monomer to oligomer | Micro |
| 15 | Petrella 2019 | Computational Causal Modeling of the Dynamic Biomarker Cascade in Alzheimer's Disease. | $A\beta$, tau, tau pathology (p-tau) | Micro |
| 16 | Proctor 2012 | A unifying hypothesis for familial and sporadic Alzheimer's disease. | $A\beta$, Tau | Macro |
| 17 | Puri 2010 | Mathematical modeling for the pathogenesis of Alzheimer's disease. | $A\beta$, microglia, astroglia | Macro |

**Table 2.** Mathematical approaches.

|  | Study | Model type | If ODE, what solver was used | Sensitivity Analysis |
| --- | --- | --- | --- | --- |
| 1 | Anastasio 2011 | Not mentioned | Maude | No |
| 2 | Bertsch 2017 | PDE | Not mentioned | No |
| 3 | Fornari 2019 | PDE, ODE and Graph | Not mentioned | No |
| 4 | Han 2020 | ODE | 5th order Runge-Kutta method | No |
| 5 | Han 2016 | Seven ordinary equations | Not mentioned | No |
| 6 | Hao 2016 | ODE and PDE | Not mentioned | Yes |
| 7 | Helal 2014 | ODE | Not mentioned | No |
| 8 | Helal 2019 | ODE | Not mentioned | No |
| 9 | Hoore 2020 | ODE | Not mentioned | Yes |
| 10 | Kuznetsov 2018 | PDE | MATLAB's BVP4C solver | No |
| 11 | Kuznetsov 2018 | PDE | Not mentioned | No |
| 12 | Lindstrom 2021 | ODE and PDE | Python | Yes |
| 13 | Liu 2015 | Not mentioned | Not mentioned | No |
| 14 | Pallitto 2001 | ODE | Not mentioned | No |
| 15 | Petrella 2019 | ODE | MATLAB | No |
| 16 | Proctor 2012 | Not mentioned | Not mentioned | No |
| 17 | Puri 2010 | ODE | Not mentioned | Yes |

## Results

### Study selection

After our search 846 studies were uploaded from PubMed to Covidence on March 29, 2022. After removing duplicates and reviewing titles and abstracts, 59 references appeared to meet our criteria. After full-text review, 41 were further excluded due to them not proposing a mechanistic model (*n* = 19); having too limited a scope (e.g., only one protein) (*n* = 12); not related to Alzheimer’s disease (*n* = 7), or being a Review or another unsuitable article form (*n* = 4) (see Fig 1), leaving 17 papers to be studied.

**Fig 1.**
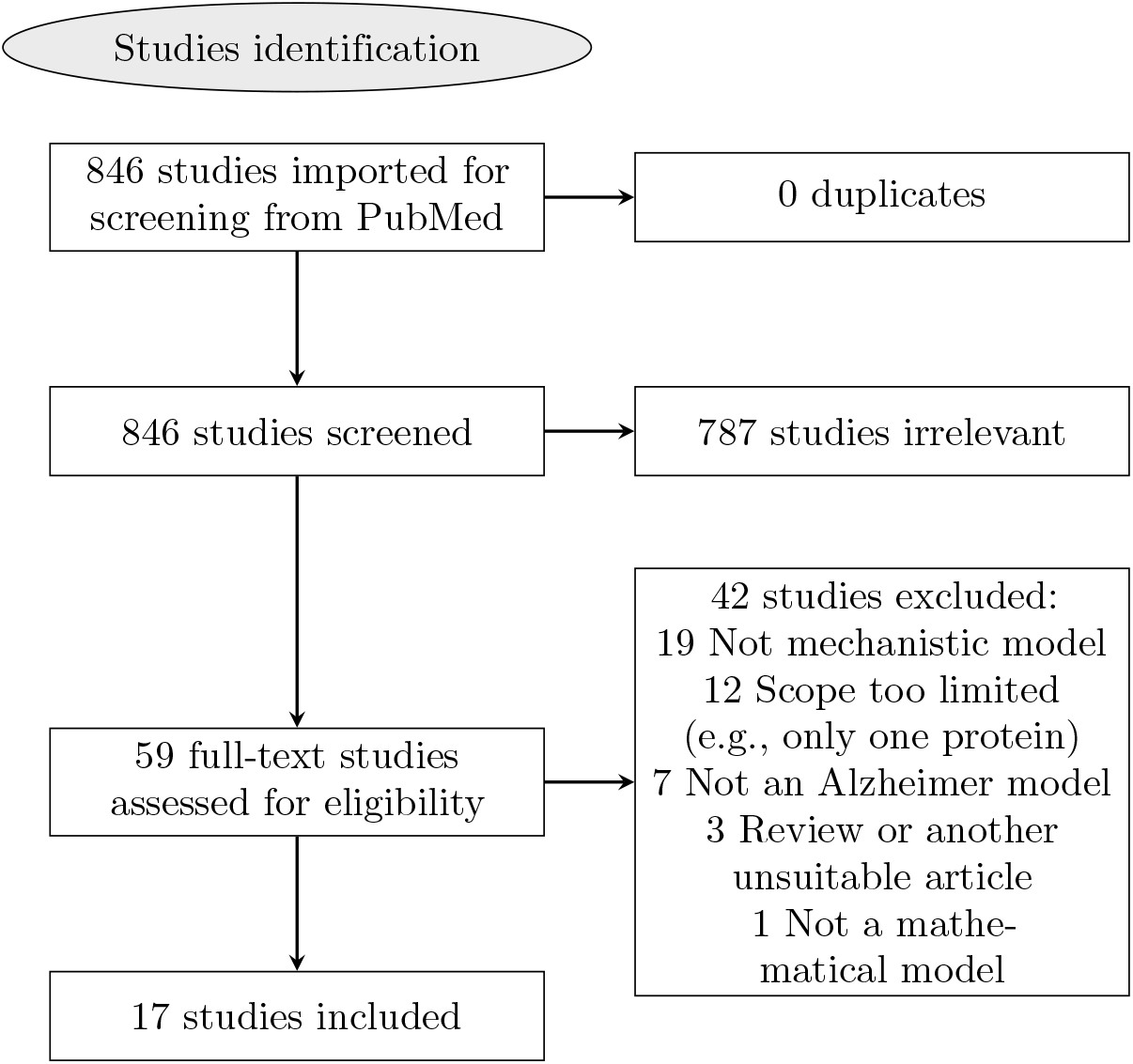
PRISMA flow chart diagram.

### Study characteristics

The 17 papers were published between 2001 and 2021. The primary authors’ countries of origin were the United States (*n* = 11) and others (Algeria, China, France, Germany, Italy, and United Kingdom).

The majority of articles modeled AD at the cellular range of the disease process, operating on a short time scale (e.g., minutes; hours), i.e., the *micro* view (12/17); while the rest of the models considered regional or brain-level processes, with longer timescales (e.g., years, decades) (the *macro* view). Most of the papers were concerned primarily with *Aβ* (*n* = 8), few modeled both *Aβ* and tau proteins (*n* = 3), and some considered more than these two factors in the model (*n* = 6) (see Table 1).

### Mathematical approaches

Mathematical approaches are summarized in Table 2. Various approaches were used, such as partial differential equations (PDE; *n* = 3), ordinary differential equations (ODE; *n* = 7), both PDE and ODE (*n* = 3), while for the remainder (*n* = 4), the type of model was not presented and only results were discussed. Of these last articles, one study used *Maude*, a language able to represent and evaluate models created using equations and rules [5]. The choice of using either ODEs or PDEs depended on how the spatial propagation of AD in the brain was considered. Models relying on PDEs considered the brain as a continuous medium with a realistic geometry, while models using ODEs either neglected the spatial component altogether or considered only a few homogenized regions.

Of the ODE models, four had components behaving over several time scales, from cellular reactions lasting minutes to disease development over years; the other six had a single time scale. The description of phenomena with characteristic time constants ranging over several order of magnitudes leads to so called *stiff* systems of ODEs. The resolution of stiff differential equations poses several numerical difficulties and the choice of an inappropriate resolution method can lead to false conclusions. In general, we found that there was not enough support or information given in the articles about which solvers and tools were used. Only four papers [8, 11, 19, 28] out of 10 mentioned which environment they use to solve their ODE system (three with MATLAB; one with Python [19]. Among these four papers, only two specifically mentioned which solvers were used. This limits our appreciation of their work.

Performing sensitivity analysis (**SA**) in mathematical models is critical to identify which parameters have an important impact on the solutions and to quantify uncertainty. SA can identify crucial model inputs (parameters and initial conditions) and quantify the impact of input uncertainty on the model’s output(s) [21]. We thus investigated whether the reviewed papers performed SA and if so, which approach was used. We found that little attention appears to have been given to this crucial subject since SA is covered but for a small number of papers (4/17).

## Discussion

### Summary of findings

The significance of AD for patients, caregivers, and society is well established. Given that there is no known cure for this disease, identifying relationships between biological factors leading to its causality is paramount, as any insight can help generate therapeutical approaches to delay or stop its progression, reducing suffering and costs [1]. Mathematical models can be extremely beneficial in this area. In this scoping review, we identified and presented findings from 17 papers with mathematical models of AD.

### Mathematical considerations

Mathematical models using continuous time, as is relevant in the case of AD, can rely either on systems of ODEs when the description of brain geometry is omitted or greatly simplified or on systems of PDEs when a continuous spatial description is performed. Either approach is *a priori* possible depending on whether one wants to emphasize or not the spatial nature of disease propagation.

A first challenge relates to the determination of parameters for these systems of equations. Given the various components involved in AD onset and progression, mathematical models will necessitate several variables and therefore be both high dimensional and nonlinear. Each interaction between variables will need to be defined using parameters describing rates and relative importance of multiple biological phenomena. Since many or most of these parameters cannot be directly measured, the objective determination of their values (or model calibration) is challenging. Therefore, the process of calibrating a mathematical model should ideally be done by comparing the model’s outputs to actual clinical, epidemiological, or experimental data [4]. Then, parameters are chosen as to minimize the difference between the model predictions and observed data. The possibility to fully characterize a model is dependent on the quantity and quality of available data [30]. In the case of AD, recent initiatives aimed at the acquisition of large publicly accessible datasets promise to increase the relevance and importance of mathematical models.

In this review, there was a wide discrepancy between the approaches taken to determine and report model parameters. Some authors used parameter values found in other modelling or experimental studies or derived them from theoretical calculations (e.g. [11, 14, 19]). However, most papers either did not explain the provenance of their parameters or their calibration process (e.g. [5, 7, 8, 10, 16, 17]), or listed parameter values without providing references or calculations (e.g. [12, 26]). Only a few papers provided experimental results to support their parameter choice (e.g. [20, 25, 28]).

Studies should provide parameter calibration and model validation approaches to increase our ability to judge whether the model reached accurate predictions [29]. When it is impossible to assess model parameters directly, their values should be determine by comparing the model’s parameters with experimental or clinical observations. In this case, optimization techniques are needed to minimize the difference between model predictions with observations [29]. Details of these techniques (and ideally the code to perform them) as optimization in highly dimensional non linear systems is far from a trivial task [29]. Depending on data type and data quality, there might be a great extent of uncertainty with respect to parameter values inferred in this way, hence the importance of specifying methodological details in order to increase reproducibility [29].

A second mathematical challenge comes from the fact that time scales ranging over several orders of magnitudes are typically involved in biomathematical models describing the onset and progression of AD. For instance, such models could include rapid protein transformation as well as a description of the disease progression over several years, as mentioned in Lindstrom et al.: “There are fast time scales for dimer dissociation (*ms*); intermediate time scales for monomer decay (*h*); and longtime scales for changes in kinetic rate constants and loss of neuronal health (*decades*)” [19]. The work by Puri et al [28] is a good example, with parameters scaling from 10^*−*5^ to 1 (1*/year*). From a mathematical standpoint, systems with components behaving over several time scales are described by stiff differential equations [23] for which numerical resolution is challenging and necessitates well-adapted specific approaches.

We found in our review that many authors did not detail which solver they used, and/or did not give sufficient information on the numerical methods they used to solve the systems of equations. To wit, there are multiple stiff ODE solvers in MATLAB (e.g. ode15s, ode23s, ode23t, and ode23tb) or Python (BDF and Radau in the SciPy library). This makes unclear whether the numerical difficulties associated with the stiffness of the model were appropriately handled. It is unfortunate that authors missed including this vital part in the description of their models, as it is necessary for reproducibility.

Finally, a third challenge in the development and analysis of high dimension mathematical models with a large numbers of parameters is related to sensitivity analysis and uncertainty quantification. Performing sensitivity analysis is essential to assess the relative importance of the contribution of each parameters [13]. Parameters having little impact on model outcome cannot be determined only by comparing model predictions to actual data [29]. Rather, several mathematical approaches can be used to perform sensitivity analysis, each providing complementary information [19, 21]. Hao et al. [11] is such an example in which the authors performed sensitivity analysis using Latin hypercube sampling to generate 2000 samples, evaluated a variation range between half and twice the parameter values, and calculated partial rank correlation coefficients and *p* − values for the density of neurons and the concentration of astrocytes at time *t* = 10 years. Another approach used by Lindstrom et al. [19] was to evaluate how tiny perturbations (1% change in a parameter value) affected model outcome, then evaluating the probability that the AD development rate differed significantly from their initial estimate. Puri et al. [28] took into consideration the parameters’ sensitivity to changes in values that are established after 20 years for a 62.5% perturbation range for each value. They reported strong, moderate, or weak effects for each parameter alongside a short explanation of their sensitivity analysis process [28]. Unfortunately, not all authors were equally forthcoming. Some (e.g. Helal et al. [13]) performed a sensitivity analysis, however did not report their method. Most others simply did not perform such an analysis. We propose as a good practice for model reporting that it should include a sensitivity analysis, complete with a description of the approach, code or pseudo-code, and numerical estimates for each parameter obtained via several simulations with different input elements focused around a nominal value.

### Internal and external validity

Determining internal and external validity is a critical but sometimes overlooked aspect of mathematical modelling. Table 3 summarizes the extent to which papers satisfied criteria of internal and external validity.

**Table 3.** Internal and external validity.

|  | Study | Given equations and initial conditions | Given parameters and their units | Sensitivity Analysis | Given code | Information on model solver | Internal validity degree | external validity |
| --- | --- | --- | --- | --- | --- | --- | --- | --- |
| 1 | Anastasio 2011 | No | No | No | No | Maude | 1/5 | No |
| 2 | Bertsch 2017 | Yes | No | No | No | No | 1/5 | No |
| 3 | Fornari 2019 | Yes | No | No | No | No | 1/5 | No |
| 4 | Han 2020 | Yes | Yes | No | No | 5th order Runge-Kutta method | 3/5 | No |
| 5 | Han 2016 | Yes | No | No | No | No | 1/5 | No |
| 6 | Hao 2016 | Yes | Yes | Yes | No | No | 3/5 | No |
| 7 | Helal 2014 | Yes | No | No | No | No | 1/5 | No |
| 8 | Helal 2019 | Yes | Yes | Yes | No | No | 3/5 | No |
| 9 | Hoore 2020 | Yes | Yes | No | No | No | 2/5 | Yes |
| 10 | Kuznetsov 2018 | Yes | No | No | No | MATLAB's BVP4C solver | 2/5 | No |
| 11 | Kuznetsov 2018 | Yes | No | No | No | No | 1/5 | No |
| 12 | Lindstrom 2021 | Yes | Yes | Yes | Yes | Python | 5/5 | Yes |
| 13 | Liu 2015 | No | No | No | No | No | 0/5 | Yes |
| 14 | Pallitto 2001 | Yes | Yes | No | No | No | 2/5 | Yes |
| 15 | Petrella 2019 | Yes | Yes | No | No | MATLAB | 3/5 | No |
| 16 | Proctor 2012 | available online | available online | No | No | No | 2/5 | No |
| 17 | Puri 2010 | Yes | Yes | Yes | No | No | 3/5 | No |

Internal validity is related to model verification and takes into account the mathematical underpinning of the model. An important aspect of internal validity is the verification of the accuracy of numerical solutions. For a mathematical model to be considered internally valid, the software and numerical methods utilized in solving the model should be specified. The code or pseudo code used should be accessible and subject to verification. Techniques like static and dynamic testing should also be used to confirm reliability, efficiency, and robustness of the code. [30]. In order to determine the extent of internal validity of the papers, we checked whether the authors provided explicit equations, initial conditions and parameter values and if so whether this was sufficiently justified. We also verified if codes or pseudo-codes were provided and if sensitivity analyses were performed.

External validity, also known as model validation, compares simulation results and model predictions to experimental data. To validate a model, experimental data from real-world experiments are used to confirm that the predictions made are accurate [30]. As can be appraised from Table 3, few articles were considered both internally and externally valid. Future reports are encouraged to provide information regarding validity as a general condition for model acceptability.

### Limitations

As a scoping review, we used a limited set of search terms (without synonyms) and only one database (PubMed). This obviously means that many studies will have been missed however, commonalities emerged out of the representative sample of studies that were reviewed. A second limitation comes from the wide scope of modeling that was proposed by the authors. While many looked at individual steps in a pathological cascade (especially that related to amyloid accumulation) few modeled the entire disease course from onset to neuronal death.

## Conclusion

The fact that there is currently no effective treatment for Alzheimer’s disease means that efforts to understand its etiology remain critical, as this knowledge is essential to devise any intervention aimed at delaying its onset. More research is therefore needed and computational models can participate in the finding of solutions. However, it is crucial to pay close attention to various concepts when creating and reporting such models. Among other things, the type of solvers used to solve ODE or PDE systems should be taken into consideration, as well as whether or not the system is stiff and if so, what strategy was used to solve the equations. Internal and external validity would be increased if equations and codes were systematically provided. Parameters and their units, the reference for each parameter, and support from experimental data are also required. Finally, a sensitivity analysis should be performed to thoroughly investigate parameters and their effects on model outputs. Such recommendations would improve our confidence in the models being proposed, and reach a stage where they can make significant testable predictions.

## Acknowledgments

S.M. is supported by a grant to S.D. from the Canadian Institutes for Health Research.

